# Dihydroartemisinin-loaded Magnetic Nanoparticles for Enhanced Chemodynamic Therapy

**DOI:** 10.1101/2019.12.20.885400

**Authors:** Shengdi Guo, Xianxian Yao, Qin Jiang, Kuang Wang, Yuanying Zhang, Haibao Peng, Jing Tang, Wuli Yang

**Author notes:** **Correspondence**: Wuli Yang or Jing Tang or Haibao Peng.

## Abstract

Recently, chemodynamic therapy (CDT) has represented a new approach for cancer treatment with low toxicity and side effects. Nonetheless, it has been a challenge to improve the therapeutic effect through increasing the amount of reactive oxygen species (ROS). Herein, we increased the amount of ROS agents in the Fenton-like reaction by loading dihydroartemisinin (DHA) which was an artemisinin (ART) derivative containing peroxide groups, into magnetic nanoparticles (MNP), thereby improving the therapeutic effect of CDT. Blank MNP were almost non-cytotoxic, whereas three MNP loading ART-based drugs, MNP-ART, MNP-DHA, and MNP-artesunate (MNP-AS), all showed significant killing effect on breast cancer cells (MCF-7 cells), in which MNP-DHA were the most potent. What’s more, the MNP-DHA showed high toxicity to drug-resistant breast cancer cells (MCF-7/ADR cells), demonstrating its ability to overcome multidrug resistance (MDR). The study revealed that MNP could produce ferrous ions under the acidic condition of tumor microenvironment, which catalyzed DHA to produce large amounts of ROS, leading to cell death. Further experiments also showed that the MNP-DHA had significant inhibitory effect on another two aggressive breast cancer cell lines (MDA-MB-231 and MDA-MB-453 cells), which indicated that the great potential of MNP-DHA for the treatment of intractable breast cancers.

## 1 Introduction

Chemodynamic therapy (CDT) is a tumor therapeutic strategy which generates abundant reactive oxygen species (ROS) in tumor sites *via* the Fenton reaction or Fenton-like reaction (Tang et al., 2019; Yan et al., 2019). Generally, specific nanomaterials produce ions as catalysts, which cleave the endoperoxide linkages in ROS agents to produce ROS (Bokare and Choi, 2014). In the classical Fenton reaction, the catalyst is ferrous ions produced under the acidic condition of tumor microenvironment and the ROS agent is the excessive hydrogen peroxide (H_2_O_2_) in cancer cells (Li et al., 2015; Chen et al., 2017). The overproduction of ROS is cytotoxic, which could damage membrane and oxidize lipids in cells, further leading to antitumor performance *via* apoptosis and/or ferroptosis (Reed and Pellecchia, 2012; Yue et al., 2018; Wan et al., 2019; Xu et al., 2019b). Owing to the fact that CDT needs to be activated by the stimulation of the tumor’s endogenous microenvironment, for example, low pH and elevated H_2_O_2_ concentration, the overproduction of ROS is almost exclusively achieved at the tumor site and consequently CDT has very low toxicity and side effects on normal tissues (Breunig et al., 2008; Chen et al., 2017; Chen et al., 2019). Compared with other treatment strategies displaying non-negligible dark toxicity, like chemotherapy, radiotherapy, photodynamic therapy, and sonodynamic therapy, CDT has the advantage that it is highly selective and specific (Osaki et al., 2011; Song et al., 2016; Cho et al., 2017; Men et al., 2018; An et al., 2019; Yang et al., 2019). However, the generation of ROS will be limited to the conditions of the tumor site, so the ideas of inducing preferential cancer cell death through exogenous ROS generating agents have gained considerable momentum.

Since the efficiency of ROS production by Fenton or Fenton-like reaction is dependent on catalysts and ROS agents, a series of studies have enhanced intracellular ROS production mainly in two aspects. On one hand, varieties of materials increasing the amount of ROS are developed from the perspective of catalysts (Zheng et al., 2017; Ma et al., 2019). Increasing the number of catalyst ions is a straightforward method to promote the efficiency of CDT. Shi group reported the facile synthesis of amorphous iron nanoparticles, which could be rapidly ionized to release Fe^2+^ ions in an acidic tumor microenvironment for CDT (Zhang et al., 2016a). Besides iron ions, many other metal ions, including Mn^2+^, Cu^2+^, and Co^2+^ ions, could also show Fenton-like activities (Ember et al., 2009; Xu et al., 2011; Bokare and Choi, 2014; Poyton et al., 2016). Due to the GSH depletion property of MnO_2_, Chen group used MnO_2_-coated mesoporous silica nanoparticles to destroy tumor cells, resulting in GSH depletion-enhanced CDT (Lin et al., 2018). On the other hand, despite the concentration of H_2_O_2_ in tumor cells is higher than normal tissues, the amount of H_2_O_2_ is still too low to achieve good therapeutic effect (Szatrowski and Nathan, 1991). Therefore, from the perspective of ROS agents, it is viable to raise the efficiency of ROS production *via* increasing the amount of ROS agents in cancer cells (Huo et al., 2017). Ge group constructed integrated multifunctional polymeric nanoparticles in which ascorbyl palmitate molecules can selectively generate H_2_O_2_ in tumor tissues, sequentially improving the therapeutic effect of CDT (Wang et al., 2018).

In addition, it is also a feasible way to load drugs whose treatment principles are based on Fenton or Fenton-like reactions into materials to increase the quantity of ROS agents. Many reports have shown that artemisinin (ART) and its derivatives, as frontline drugs against malarial infections, achieve antimalarial effects by Fenton-like reaction, the specific process of which is that under the catalysis of ferrous heme the weak endoperoxide linkages (R-O-O-R’) in drugs break resulting in the formation of toxic ROS (Olliaro et al., 2001; Krishna et al., 2004; Golenser et al., 2006; Tu, 2011). Currently, ART and its derivatives have also been used as tumor therapeutic agents for cancers *via* CDT (Wang et al., 2016; Yao et al., 2018; Sun et al., 2019; Wang et al., 2019b). What’s more, it has been found that ART and its derivatives showed sensitivity against multidrug resistance (MDR) cancer cells, as that some common ART-based drugs were not transported by P-glycoprotein (P-gp), which mediates cellular MDR by actively pumping antitumor drugs outside the cancer cells (Kruh and Belinsky, 2003; Szakacs et al., 2006; Prasad et al., 2012; Zhong et al., 2016; Wang et al., 2019c). Therefore, ART and its derivatives exhibit the potential to overcome tumor MDR.

In this work, as shown in **Scheme 1**, ART and its two derivatives, dihydroartemisinin (DHA) and artesunate (AS), were loaded into magnetite nanoparticles (MNP) respectively used for CDT enhancement. After loading these drugs, the non-cytotoxic MNP showed high toxicity to breast cancer cells. Subsequently, dihydroartemisinin-loaded magnetic nanoparticles (MNP-DHA) with the best inhibitory effect exhibited the ability to effectively kill MCF-7/ADR cancer cells, and the mechanism of MNP-DHA achieving therapeutic effect was investigated. Further experiments indicated that MNP-DHA possessed excellent inhibition ability for other intractable breast cancer cells and had a good application prospect.

**SCHEME 1.**
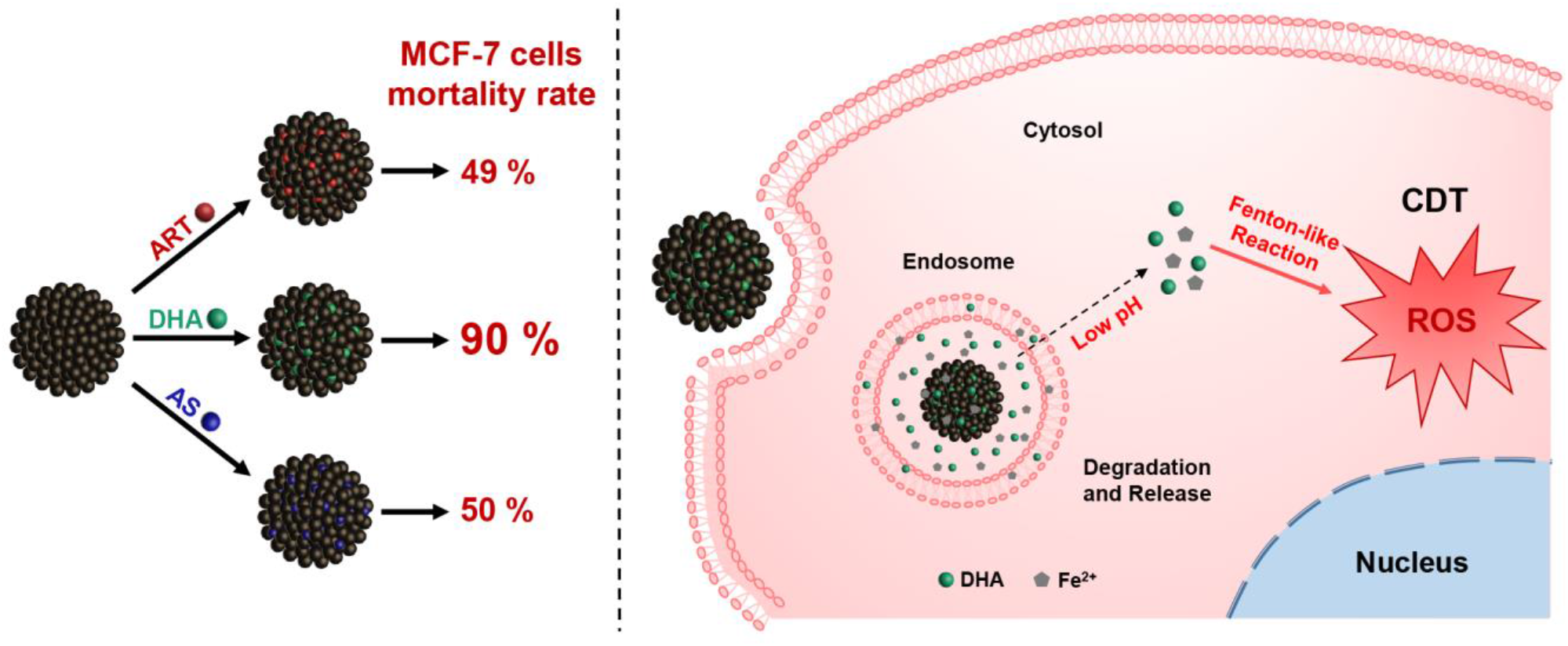
Schematic illustration of dihydroartemisinin-loaded magnetic nanoparticles for enhanced chemodynamic therapy.

## 2 Materials and methods

### 2.1 Materials

Iron (III) chloride hexahydrate (FeCl_3_·6H_2_O), sodium acetate anhydrous (NaOAc), trisodium citrate dihydrate (Na_3_Cit·2H_2_O), ethanol, sodium hydroxide (NaOH), and dimethyl sulfoxide (DMSO) were purchased from Shanghai Chemical Reagents Company. Doxorubicin hydrochloride (DOX), artemisinin (ART), dihydroartemisinin (DHA), artesunate (AS), 1,3-diphenylisobenzofuran (DPBF), sodium dihydngen phosphate anhydrous (NaH_2_PO_4_) and sodium phosphate dibasic anhydrous (Na_2_HPO_4_) were purchased from Shanghai Aladdin Chemistry Company. 2’,7’-dichlorofluorescein diacetate (DCFH-DA) and cell Counting Kit-8 (CCK-8) were purchased from Keygen Biotech Company (Nanjing, China). FerroOrange was purchased from Dojindo Molecular Technologies Company. Roswell Park Memorial Institute-1640 (RPMI-1640) medium, Dulbecco’s modified Eagle’s (DMEM) medium, penicillin/streptomycin solution, fetal bovine serum (FBS), and trypsin-ethylene diamine tetraacetic acid (Trypsin-EDTA, 0.05 %) were purchased from Gibco BRL (Grand Island, NY). The water used in the experiment was deionized water.

### 2.2 Characterization

The morphology of nanoparticles was tested by a Tecnai G2 20 TWIN transmission electron microscope (TEM) at an accelerating voltage of 200 kV and a Zeiss Ultra 55 field emission scanning electron microscope (FESEM) equipped with a fieldemission gun operated at 5 kV. Magnetic characterization curves were measured by a Quantum vibrating sample magnetometer (VSM) at 300 K. Dynamic light scattering (DLS) data, including the size, zeta potential and light scattering intensity of the nanoparticles were measured at 25 °C on a Zetasizer Nano ZS90 analyzer (Malvern Instrument Ltd). Fourier transform infrared (FT-IR) spectra were obtained *via* a FT-IR spectrometer (Thermofisher Nicolet 6700). Ultraviolet spectrophotometer (UV-Vis) spectra were recorded at 25 °C on a Perkin-Elmer Lambda 750 spectrophotometer. The concentration of metal ions was obtained on a P-4010 inductively coupled plasma-atomic emission spectrometry (ICP-AES). Confocal laser scanning microscopy (CLSM) images were acquired using a Nikon C2+ laser scanning confocal microscope. Flow cytometry analysis was operated on a flow cytometer (Beckman Coulter Gallios) at 37 °C.

### 2.3 Synthesis of Magnetic Nanoparticles

Magnetic nanoparticles (MNP) were prepared *via* a modified solvothermal reaction (Wang et al., 2019a). FeCl_3_·6H_2_O (1.8 g), Na_3_Cit·2H_2_O (1.2 g) and NaOAc (4.8 g) were dissolved in 88 mL ethylene glycol with sonicated in an ultrasonic bath for 10 minutes, then the mixture was stirred vigorously for 30 minutes. The resulting solution was then transferred into a autoclave, which was sealed and heated for 12 h at 200 °C. After cooling down to room temperature, separated by a magnet, the product was washed alternately with ethanol and deionized water for 3 times, then redispersed in water for subsequent use.

### 2.4 Preparation and Release Study of Drug-loaded MNP *in Vitro*

Three drugs were loaded into MNP, including ART, DHA, AS, respectively. 6 mg of MNP were added into 2 mL of deionized water and then sonicated for 5 min to form a homogeneous dispersion. Then 1.5 mg of ART dissolved in 1 mL of ethanol was added to the dispersion and the dispersion was shaken up for 24 h at room temperature. Subsequently, liquid of the dispersion was removed by rotary evaporation at 40 °C. The product was washed with water for 3 times *via* a magnet and then collected for further use. After treating with NaOH-containing ethanol solution at 50 °C for 30 minutes, the unloaded ART in the collected supernatant was converted to a UV active compound and detected by a UV-visible spectrometry at an excitation wavelength of 292 nm. According to the following formulation, the drug loading contents (LC) were calculated: LC (%) = (the drug loaded in MNP weight) / (total nanoparticles weight) × 100 %.

The methods of loading DHA and AS into MNP were similar to the above method, except the mass ratio of MNP and the drug, and the volume ratio of water and ethanol. When loading DHA into MNP, 10 mg of MNP were added into 2 mL of deionized water and then 3 mg of DHA dissolved in 2 mL of ethanol was added to the dispersion. When loading AS into MNP, 10 mg of MNP were added into 4.95 mL of deionized water and then 3 mg of AS dissolved in 0.05 mL of ethanol was added to the dispersion. Furthermore, the method of converting drugs to UV active compounds was different between different drugs. In order to be measured at the wavelength of 238 nm, DHA was treated with ethanol solution containing NaOH at 60 °C for 30 min and AS was treated with NaOH solution (0.1 M) at 83 °C for 1 hour. The stability of drug-loaded nanoparticles in phosphate buffer saline (PBS, pH 7.4) was detected *via* monitoring the hydrodynamic size and polydispersity index (PDI) by DLS.

The drug release behaviors were studied *via* an incubator shaker at 37 °C. Sealed in a 1.4 × 10^4^ Dalton dialysis bag, 2 mL of drug-loaded MNP were immersed into 200 mL of PBS (pH 7.4) and incubated under oscillation. At predetermined time intervals, 2 mL of release solution was withdrawn and replaced by an equal volume of fresh buffer. Through UV-visible spectrometry, the concentration of drug released from nanoparticles was obtained. Cumulative drug release was calculated as a percentage of the total drug loaded in MNP and plotted over time. All measurements were performed three times.

### 2.5 Cell Culture

Human embryonic kidney cell line (HEK-293T cells, normal cells), human breast cancer cell line (MCF-7, MDA-MB-231, and MDA-MB-453 cells, tumor cells), and human breast drug-resistant cancer cell line (MCF-7/ADR cells, tumor cells) were purchased from Chinese Science Academy. HEK-293T, MCF-7, and MDA-MB-231 cells were cultured in DMEM supplemented with 10 % (v/v) FBS and 1 % antibiotics (penicillin/streptomycin, 100 U/mL). MDA-MB-453 and MCF-7/ADR cells were cultured in RPMI-1640 containing 10 % (v/v) FBS, 1 % antibiotics (penicillin/streptomycin, 100 U/mL) and DOX (0.5 μg/mL). Cells were incubated in an atmosphere of 5 % CO_2_ at 37 °C.

### 2.6 Cytotoxicity Assays

The cytotoxicity of nanoparticles was tested on cells using a standard CCK-8 assay (Jiang et al., 2017). Cells were incubated in 96 pore plates at an initial density of 1 × 10^4^/well for 24 h at 37 °C and under 5 % CO_2_ atmosphere. Then different concentrations of MNP, drugs and drug-loaded MNP (100 μL/well) dispersions were added in each well and coincubated with cells for 24 h, respectively. At last, CCK-8/culture medium (10 μL/100 μL) was added into each well for another 1 h incubation. The absorbance at 450 nm of each well was measured using a BioTek enzyme-linked immunosorbent assay reader. All measurements were repeated in triplicate.

### 2.7 Acid-responsive Behaviors

To investigate the acid degradation performance of MNP, the concentrations of iron ions generated *via* MNP at PBS (pH 7.4 and 5.0) were measured by an inductively coupled plasma spectrometer (ICP). MNP (200 μg/mL) were sealed in a 1.4 × 10^4^ Dalton dialysis bag and incubated in 200 mL of PBS (pH 7.4 and 5.0) at 37 °C under oscillation, respectively. At different time points, 2 mL of release solution was removed and replaced with an equal volume of fresh solution. The cumulative release of iron ions was calculated as the percentage of total iron ions in the same mass MNP and plotted with time. Each measurement was repeated three times.

### 2.8 Detection of Cellular Fe^2+^ ions Generation

To clarify Fe^2+^ ions generation *via* the nanoparticles in cells, CLSM measurement was performed. MCF-7/ADR cells were seeded in confocal dishes at the density of 1 × 10^5^ cells/mL, cultured for 24 h, and then MNP, DHA, MNP-DHA dispersions (200 μg/mL) were added into dishes, respectively. Meanwhile, a dish without adding samples was prepared as a control group. After incubated for 6 h, the culture medium was removed and cells were washed with PBS three times. Then FerroOrange (1 μM, an intracellular Fe^2+^ ions probe, Ex: 543 nm, Em: 580 nm) dispersed in serum-free medium was added to the cells, and cells were incubated for 30 min in a 37 °C incubator equilibrated with 95 % air and 5 % CO_2_. Finally, the fluorescence images of cells were captured using a C2+ confocal microscope.

### 2.9 Detection of ROS Generation *in Vitro*

In order to measure the generation of ROS, DPBF was selected as the ROS trapper, which can be oxidized by ROS resulting in fluorescence quenching (Ding et al., 2018). Typically, DPBF (10 μM), FeSO_4_·7H_2_O (100 μM) and DHA (100 μM) were dissolved in ethanol quickly, and the above mixture was measured by the UV-vis spectrophotometer for 0, 2, 5, 10, 20, 30, 60, 90 and 120 min at the wavelength of 410 nm, respectively.

The production of ROS in MCF-7 and MCF-7/ADR cells was detected by CLSM and flow cytometry (Yang et al., 2019). DCFH-DA, as a ROS probe, was used to assess intracellular ROS generation ability. Cells were seeded in confocal dishes at a density of 1 × 10^5^ cells/mL and incubated for 24 h to allow cell attachment. Then cells were incubated with different materials respectively and the plate without adding samples was as a control group. After incubated for 6 h, the culture medium was removed and cells stained with 1 mL of DCFH-DA (10 μM) dissolved in PBS at 37 °C for 30 min. Afterward, PBS containing DCFH-DA was removed and cells were rinsed three times with fresh PBS. The fluorescence images of cells were captured using a C2+ confocal microscope.

Besides, using a flow cytometer ROS production was quantitatively measured. Cells were seeded onto a 6-well plate at a density of 1 × 10^5^ cells/mL and treated as the similar steps above to be dyed. Then cells were digested and transferred into centrifuge tubes. Cells were separated *via* centrifugation for 5 min at 1000 rpm and redispersed in PBS (0.5 mL). The fluorescence intensity of DCF was tested by the flow cytometry.

## 3 Results and Discussion

### 3.1 Preparation and Characterization of Drug-loaded MNP

The synthesis method of MNP was slightly modified based on the published solvothermal method (Deng et al., 2005). The detailed morphological and structural features of MNP were examined by TEM, demonstrating the rough surface and the uniform morphology with the particle size of ~180 nm (**Figure 1A**). Meanwhile, FESEM images also showed the spherical structure of MNP (**Figure 1B**). In addition, the magnetic hysteresis curves showed no evident remanence and coercivity, suggesting superparamagnetic property of MNP (**Figure 1C**). The inset photo that MNP were separated *via* a magnet also revealed MNP had very good magnetism.

**FIGURE 1.**
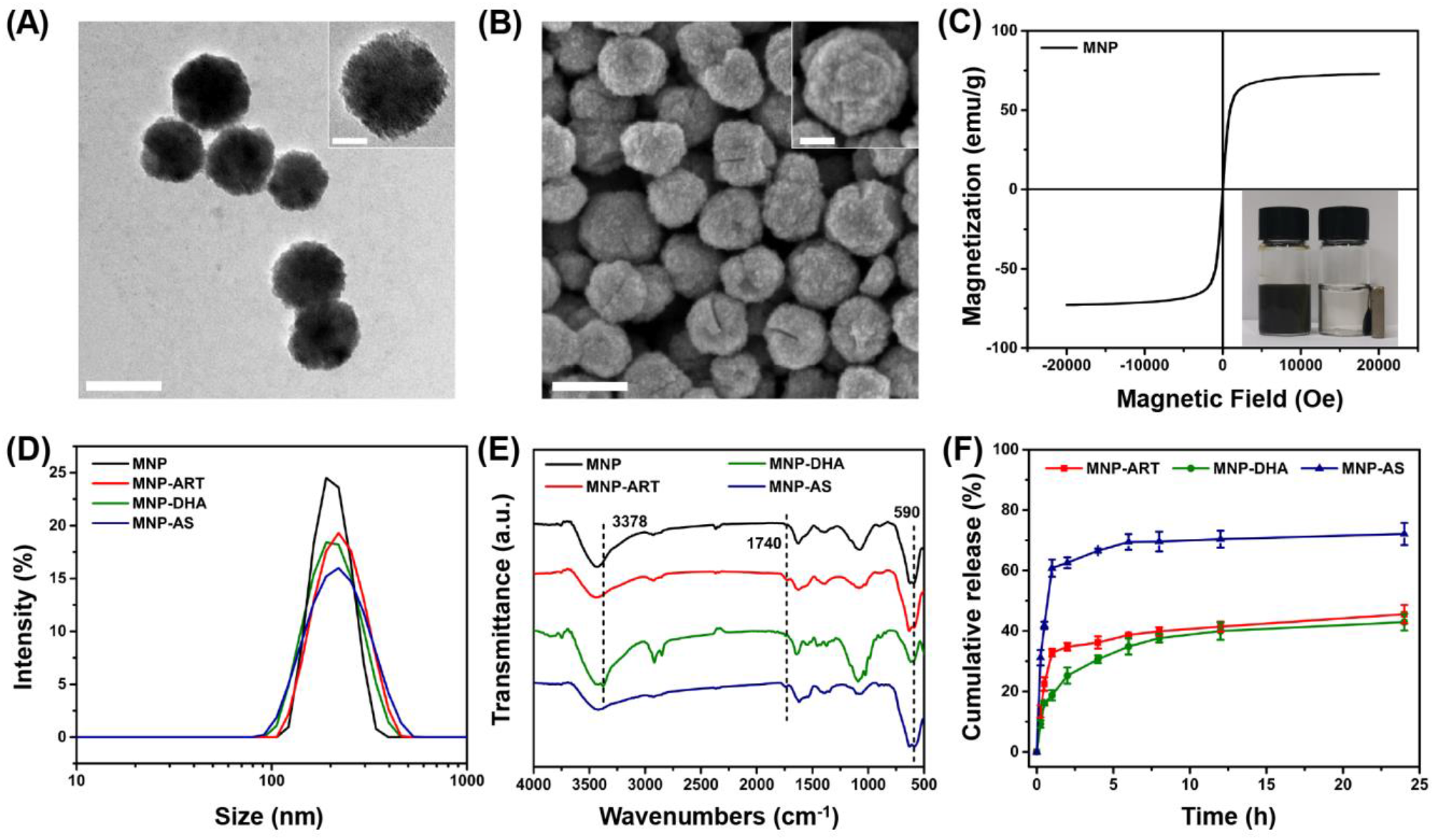
TEM images of **(A)** MNP. FESEM images of **(B)** MNP. The scale bars represent 200 nm and the scale bars of insets are 50 nm. **(C)** Magnetic hysteresis curves of MNP. **(D)** DLS curves of MNP, MNP-ART, MNP-DHA, and MNP-AS in PBS (pH 7.4). **(E)** FT-IR spectra of MNP, MNP-ART, MNP-DHA, and MNP-AS. **(F)** Cumulative drug release from MNP-ART, MNP-DHA, and MNP-AS in PBS (pH 7.4).

As shown in **Figure 1D** (and **Table S1)**, the hydrodynamic diameter (Dh) of MNP was 200 nm with a narrow PDI of 0.013. After loading drugs, including ART, DHA and AS, the average sizes of MNP-ART, MNP-DHA and MNP-AS were 212, 204 and 204 nm, and the PDI were 0.065, 0.026 and 0.092, respectively, which implied that the load of drugs didn’t affect the stability of nanoparticles. Furthermore, the particle size, as shown by DLS, was larger than that shown by TEM and SEM, which was probably due to the interaction between nanoparticles and surrounding water molecules.

The FT-IR spectra demonstrated the successful loading of drugs (**Figure 1E** and **Figure S1**). The characteristic peak at 590 cm^−1^ was attributed to Fe-O bond (Sanati et al., 2019). After ART loading, the spectrum of the MNP-ART exhibited new band in the 1740 cm^−1^ region, which belongs to C=O in δ-lactone of ART. In the same way, the absorption peaks at 3378 and 1740 cm^−1^ belong to O−H of DHA and C=O of AS, respectively (Ding et al., 2018; Kumar et al., 2019). The loading ratios of the three drugs were further measured by the UV-vis spectra. According to the standard curves of three drugs (**Figure S2**), the LC could be calculated that ART, DHA and AS were loaded in MNP with contents of 15.3 %, 15.3 % and 15.7 %, respectively. By the way, the LC of three drugs were all very close to 15 %, which was deliberately controlled *via* adjusting the mass ratio of MNP to the drug, and with the similar drug LC, latter experiments could be more comparable.

In order to understand the drug release behavior, the drug release profiles of drug-loaded MNP were investigated. As shown in **Figure 1F**, the cumulative release of ART was about 45.5 % and DHA was about 42.9 % over 24 h, which confirmed that the capacities of MNP to hold ART and DHA in physiological environment were similar. Actually, the solubility of DHA was slightly lower than ART, so during the first 2 hours of the release process, ART exhibited a distinct rapid release behavior, which DHA didn’t (Wang et al., 2007; Ansari et al., 2011). In addition, the cumulative release of AS reached 72.1 % over 24 h, indicating that AS was more hydrophilic than ART and DHA, which was consistent with the reported work (Xu et al., 2019a).

### 3.2 *In Vitro* Biocompatibility and Cytotoxicity Assays

The cytotoxicity of nanoparticles to different cells was assessed using CCK-8 assays (Jiang et al., 2017). As shown in **Figure S3**, after incubation with blank MNP for 24 h, there was no obvious toxic effect on HEK-293T cells, and cell viability retained above 90 % even with a high concentration up to 200 μg/mL, which indicated good biocompatibility of blank MNP.

To evaluate the cytotoxicity of ART and its derivatives to cancer cells, MCF-7 cells were incubated with blank MNP, the drugs and the drug-loaded MNP for 24 h, respectively. As shown in **Figure 2A, 2B** and **2C**, all CCK-8 assays displayed dose dependent cell viability. Cells treated by blank MNP still remained high viability at the concentration of 200 μg/mL. From the results of free-drug groups, the inhibitory effects of ART and AS to cancer cells were also not good enough at various concentrations. However, simultaneous delivery of drugs and MNP into cancer cells all exhibited sharply enhanced cytotoxicity. For instance, in the blank MNP group, the cell viability decreased by only 4 % at a concentration of 100 μg/mL, and at free ART, DHA, and AS concentrations of 18 μg/mL, the cell viability decreased by 4 %, 39 %, and 14 %, respectively, while in the corresponding concentrations of MNP-ART, MNP-DHA, and MNP-AS groups, the cell viability was reduced by approximately 49 %, 90 %, and 50 %, respectively, which is far greater than the sum of cell viability reduced by the two agents alone. This finding showed that ART and its derivatives had a particularly significant enhancement to MNP of inhibitory effects on cell viability, even exceeding the killing effect of the agent itself.

**FIGURE 2.**
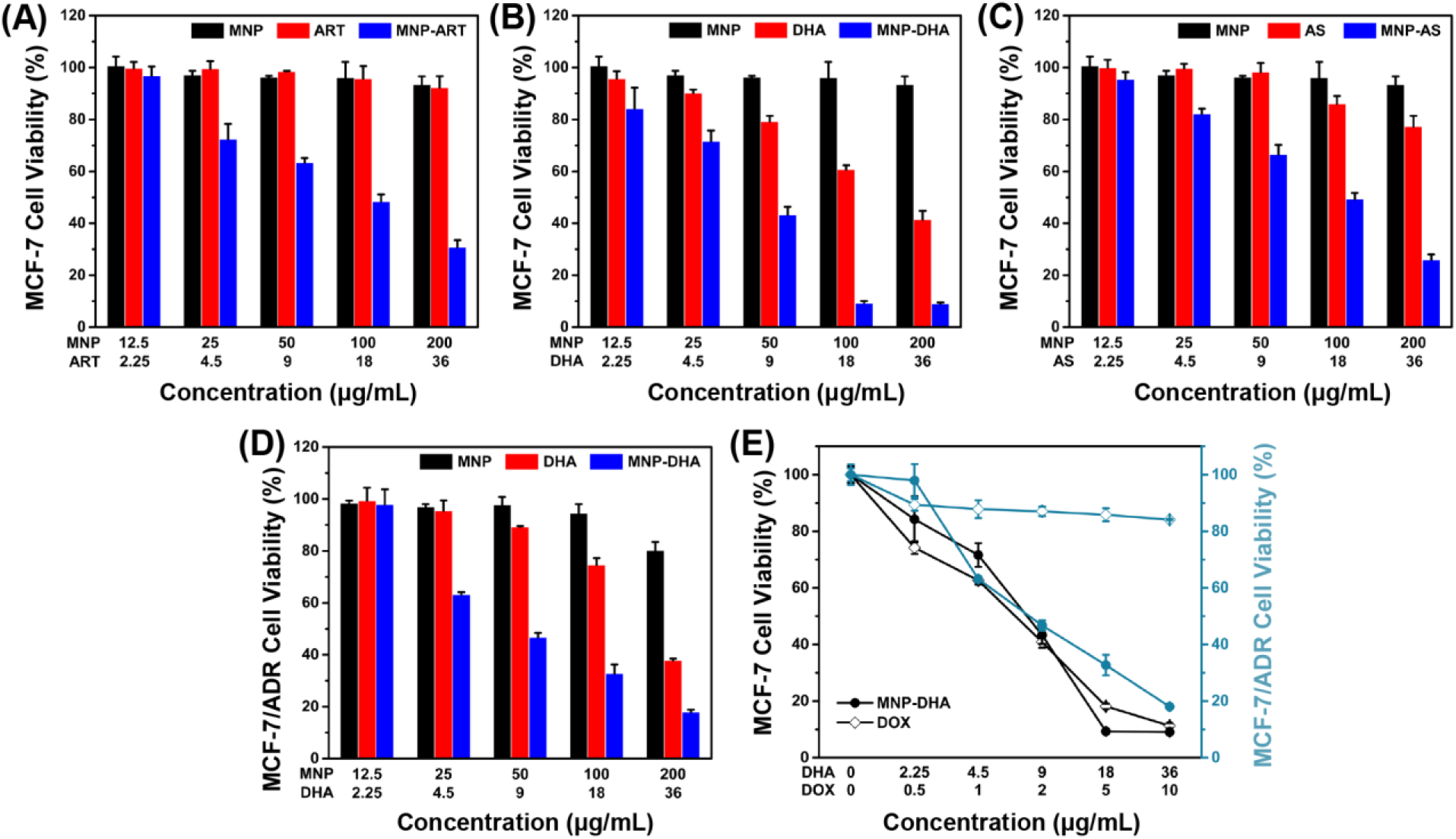
MCF-7 cell viability after incubated with MNP, free drugs, and MNP-drug dispersions at different concentrations for 24 h, respectively: **(A)** ART, **(B)** DHA, **(C)** AS. **(D)** MCF-7/ADR cell viability after incubated with MNP, free DHA, and MNP-DHA suspensions at different concentrations for 24 h. **(E)** A comparison of the inhibitory effect of MCF-7 and MCF-7/ADR cells treated with free DOX and MNP-DHA.

The MNP-DHA, which had the best effect on inhibiting cancer cell viability in three MNP loading ART-based drugs, was selected for subsequent experiments. After calculation, the half inhibitory concentration (IC_50_) of free DHA was 26.10 μg/mL, which was significantly reduced after loading into MNP, changing to 7.76 μg/mL. It was shown that MNP-DHA had a better effect on killing cancer cells than free DHA, which meant the enhancement effects of materials and drugs is mutual, and further demonstrated that the combined use of DHA and MNP was an excellent strategy for enhancing killing cells effects.

According to previous reports, ART and its derivatives were sensitive to drug-resistant tumor cells, so we tried to use MNP-DHA to carry out cytotoxicity experiments on MCF-7/ADR cell lines (Zhong et al., 2016; Hu et al., 2019). As shown in **Figure 2D**, only 33 % cell viability was obtained after treatment by MNP-DHA at a concentration of 100 μg/mL. The results showed that MNP-DHA also had a great killing effect to MCF-7/ADR cells. To compare the therapeutic effects on drug-resistant cancer cells between MNP-DHA and DOX, the cytotoxicity of DOX on the MCF-7 and MCF-7/ADR cell lines was evaluated. After treatment of cells with free DOX for 24 h, MCF-7 cell viability decreased rapidly, while the viability of MCF-7/ADR cells showed little change (**Figure 2E**). Whereas, whether MCF-7 or MCF-7/ADR cells, their survival rate became very low after treatment with MNP-DHA for 24 h. Consistent with published studies, the results showed that DHA wasn’t a P-gp substrate, as a consequence, DHA could bypass P-gp mediated MDR(Crowe et al., 2006; Wang et al., 2019c). This finding demonstrated that the proposed MNP-DHA could overcome the MDR of MCF-7/ADR cells and induce high cytotoxicity.

### 3.3 *In Vitro* Study of Fe^2+^ ions Generation

The participation of a large number of ferrous ions was essential for the high efficiency of CDT, so it was necessary to evaluate the dissolving process of MNP in an acidic environment (PBS, pH 5.0), which simulated the acidic condition in the tumor microenvironment (Breunig et al., 2008; Hao et al., 2010; Wang et al., 2016; Zhang et al., 2016b). The acid degradation experiments were carried out in PBS of different acidity (pH 7.4 and 5.0). Certified by the ICP-AES, the released iron ions increased with continuously degradation of MNP and as the pH value of PBS decreased, MNP exhibited more severe degradation. After 12 h, the Fe concentration in the pH 7.4 buffer solution was only 1.61 μg/mL, while the concentration of iron ions in the pH 5.0 buffer solution reached up to 10.45 μg/mL (**Figure 3A**), implying that MNP could be degraded into abundant iron ions in the microenvironment of tumors.

**FIGURE 3.**
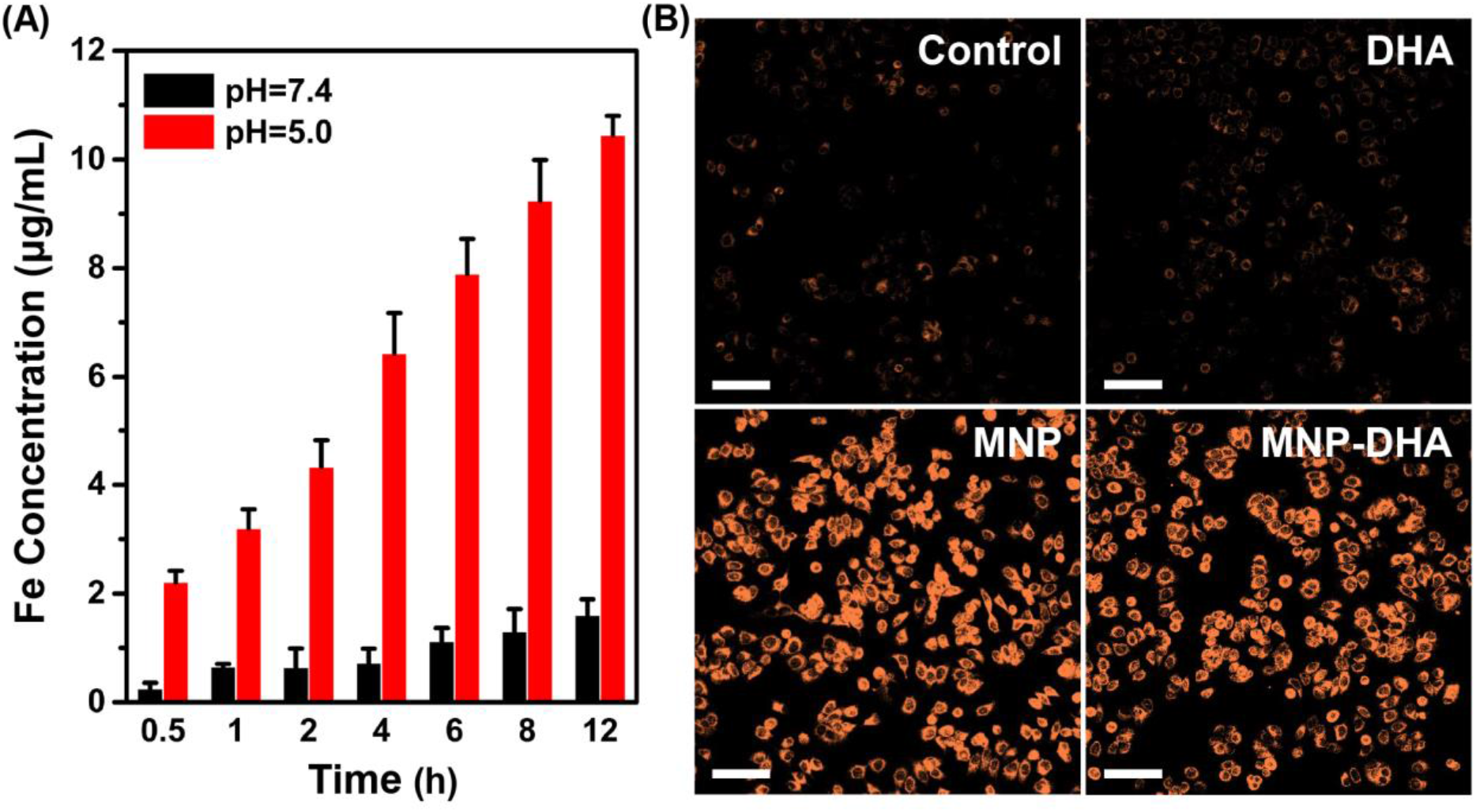
**(A)** The quantitative analysis of iron ions released from pH-sensitive MNP at different pH (7.4 and 5.0) environment. **(B)** CLSM images of MCF-7/ADR cells collected to visualize the intracellular Fe^2+^ ions generation using the Fe^2+^ ions fluorescent probe Ferrorange. The scale bars are 100 μm.

The generation of Fe^2+^ ions was corroborated using a Fe^2+^ ions probe known as Ferrorange, which could react with Fe^2+^ ions to produce a bright fluorescent substance. Compared with the control and free DHA groups, the cells treated with MNP and MNP-DHA emitted a much stronger orange fluorescence (**Figure 3B**), indicating an enormous amount of Fe^2+^ ions generated *via* MNP.

### 3.4 *In Vitro* CDT Mechanism of MNP-DHA

It was well-known that endoperoxide linkages could be cleaved with ferrous ions to generate ROS *via* a Fenton-like route, which further caused apoptosis or ferroptosis of cells (Efferth et al., 2004; Ooko et al., 2015). To understand the enhanced mechanism of DHA to CDT, an assessment of the ROS generation ability produced by the reaction of DHA with Fe^2+^ ions was investigated first. A classical ROS trapper, DPBF, was used to measure ROS generation. As the generation of ROS increased, the absorbance of DPBF decreased (Ding et al., 2018). As shown in **Figure 4A**, at the beginning of the reaction, DPBF solution had a strong absorption at 412 nm. With the reaction time increasing, the absorbance of DPBF gradually decreased, indicating that ROS was produced gradually through the interaction of DHA and Fe^2+^ ions over time.

**FIGURE 4.**
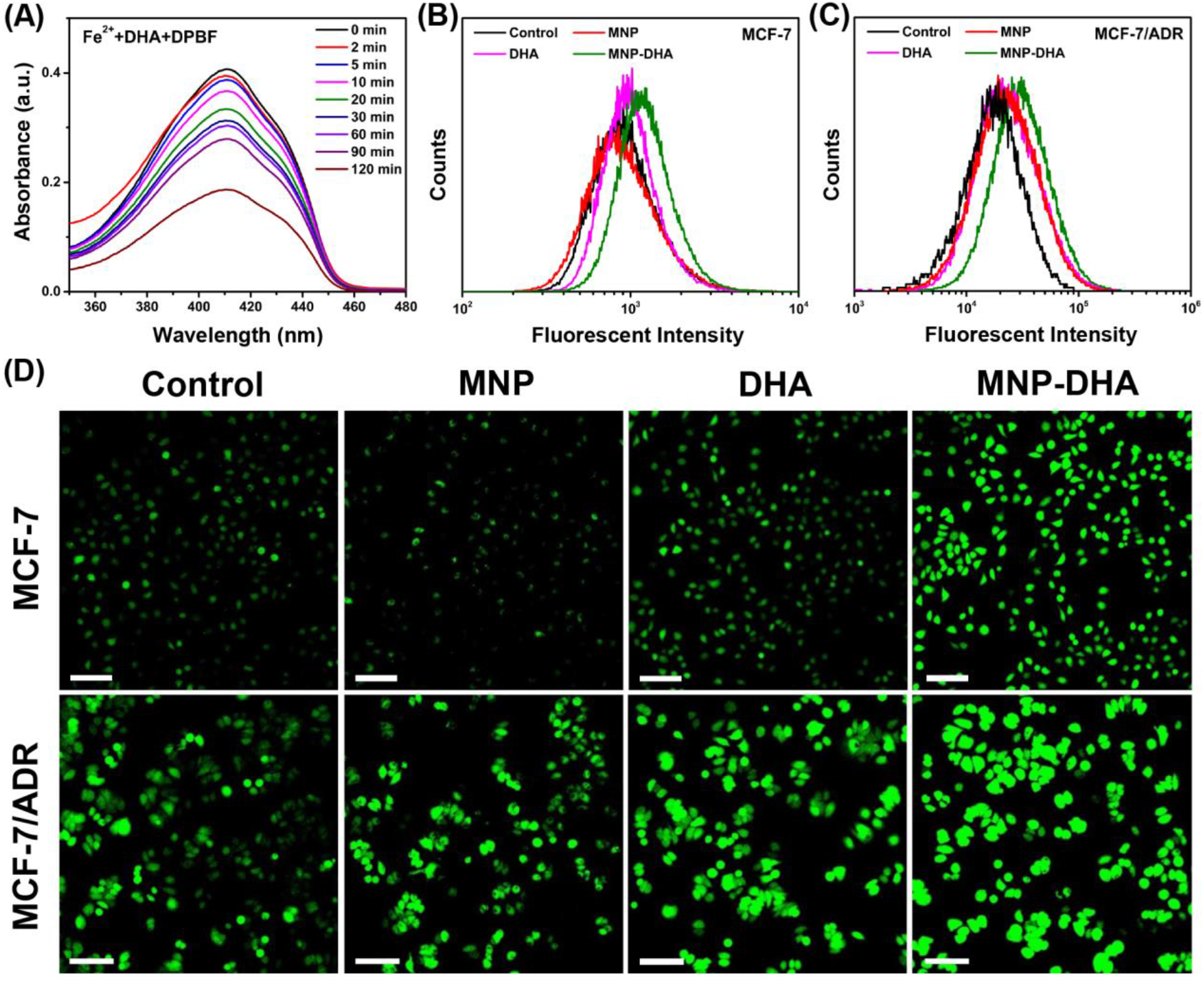
**(A)** The absorption spectra of DPBF at the presence of DHA and Fe^2+^ ions at different time. Flow cytometry analyses of ROS generation in **(B)** MCF-7 and **(C)** MCF-7/ADR cells detected by DCFH-DA. **(D)** CLSM images of MCF-7/ADR cells treated under different conditions to evaluate ROS production based on DCF fluorescence intensity using the fluorescent probe DCFH-DA. The scale bars are 100 μm.

Afterwards, we compared the ROS yielding ability of different groups by means of flow cytometry and CLSM, including control, blank MNP, free DHA, and MNP-DHA group. A fluorescent probe DCFH-DA was chosen to test intracellular ROS generation, which enable to produce fluorescent 2’,7’-dichlorofluorescein (DCF) under the combined actions of cellular esterase and ROS (Yuan et al., 2014). The quantitative fluorescence analysis was measured by flow cytometry (**Figure 4B** and **4C**). Incubated with or without MNP, the MCF-7 and MCF-7/ADR cells showed no significant difference in the fluorescence intensity of DCF, due to the fact that the concentration of H_2_O_2_ in cells was not enough to produce a large amount of ROS with ferrous ions. After incubation with free DHA, the fluorescence intensity of the cells became a little higher, on account of the reaction of naturally existed Fe^2+^ ions with DHA. After treatment with MNP-DHA, a significant enhancement of DCF fluorescence in both MCF-7 and MCF-7/ADR cells was clearly observed, owing to the ROS generation from Fe^2+^ ions and DHA brought by DHA-loaded nanoparticles. The experiments suggested that more intracellular ROS were produced after treated by MNP-DHA.

The results of fluorescence imaging agreed well with flow cytometry. As shown in **Figure 4D**, whether the cell line used in the experiments was MCF-7 or MCF-7/ADR, the fluorescence observed in control and MNP group was faintest. The fluorescence slightly increased in free DHA group, indicating that moderately amount of ROS was generated. In the MNP-DHA group, the fluorescence was greatly enhanced, which was the strongest of the four groups. Therefore, the results verified that the effect of DHA from MNP-DHA on enhancing the production efficiency of intracellular ROS was very significant.

### 3.5 Cytotoxicity Assays of Other Breast Cancer Cells

In consideration of the high cytotoxicity of MNP-DHA, we tried to use this combination to conduct toxicity experiments on other canonical lethal breast cancer cell lines that were triple negative (MDA-MB-231) and human epidermal growth factor receptor (HER2) overexpressing (MDA-MB-453) (Neve et al., 2006; Lee et al., 2012). Triple-negative breast cancer, defined by the lack of estrogen receptor, progesterone receptor and HER2, frequently developed resistance to chemotherapy over long-term treatment (Kim et al., 2018; Raninga et al., 2020). HER2 was overexpressed in 25-30 % of breast cancers which was a considerable proportion, and patients with breast cancers that overexpress HER2 had much lower overall survival and disease-free survival due to high metastasis (Baselga et al., 1998; Slamon et al., 2001; Büyükköroğlu et al., 2016). As a consequence, it was of great significance to develop novel therapies for these tumors. As shown in **Figure 5**, after mixing with 100 μg/mL of MNP-DHA for 24 h, the viability of MDA-MB-231 cells decreased to 21 % and the viability of MDA-MB-453 cells reduced to 19 %. This finding made it possible to treat other types of refractory breast cancers *via* MNP-DHA, nonetheless the specific mechanism needed further research.

**FIGURE 5.**
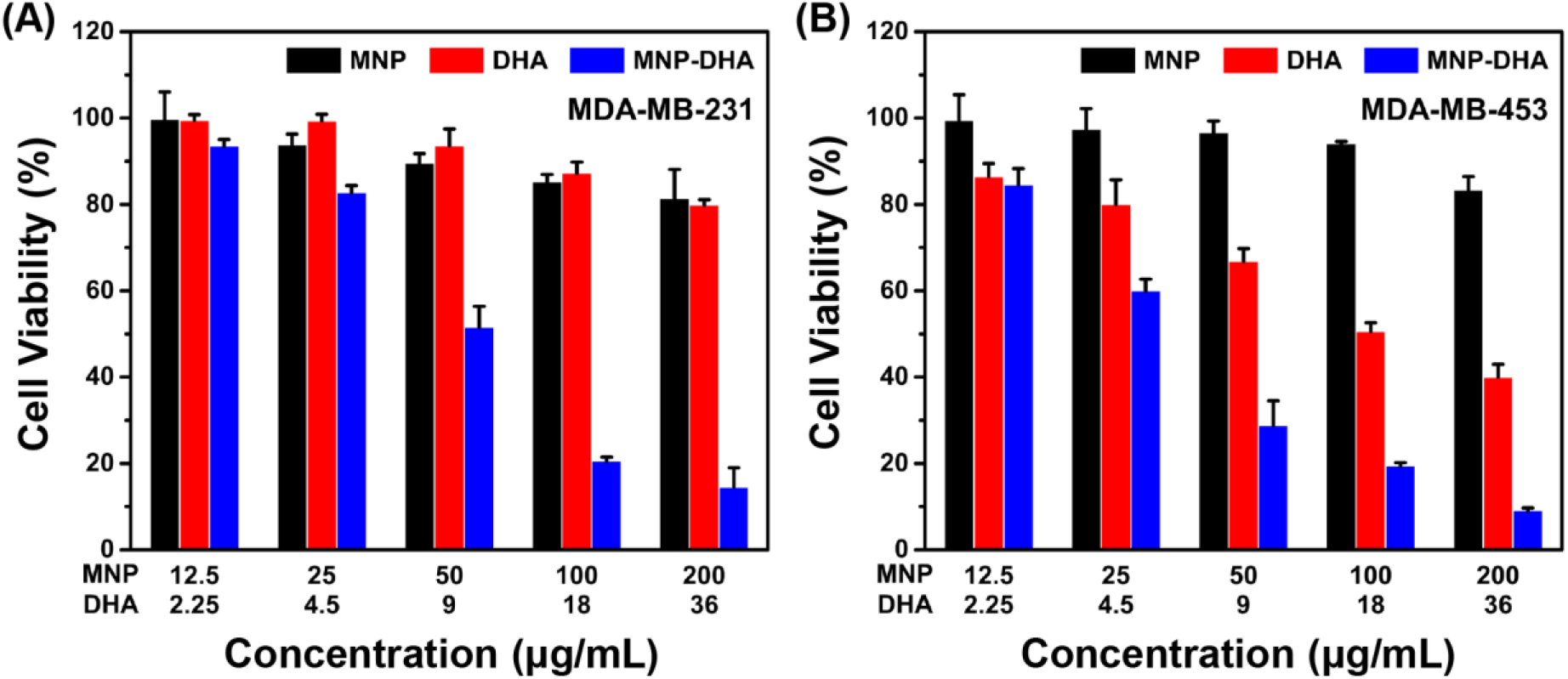
Cell viability of **(A)** MDA-MB-231, and **(B)** MDA-MB-453 after 24 h incubation with MNP, free DHA, and MNP-DHA suspensions at different concentrations.

## 4 Conclusion

In summary, we successfully improved the therapeutic effect of CDT *via* loading the drugs containing peroxide groups into MNP. Among three MNP loading ART-based drugs, MNP-DHA had the strongest inhibitory effect on breast cancer cells. MNP-DHA were capable of specifically performing the Fenton-like reaction in the tumor microenvironment, thereby producing a large amount of ROS to kill tumor cells. In addition, MNP-DHA could overcome the P-gp mediated tumor MDR and could be used to treat other aggressive breast tumors. Altogether, the proposed nanoparticles may provide an effective solution for improving the efficacy of CDT treatment and have a good prospect in the treatment of aggressive breast cancers.

## Supporting information

jing tang

## 5 Author Contributions

S.G. and W.Y. designed the research. S.G., K.W. and Y. Z. conducted the experiments. S.G., X.Y. and Q. J. analyzed the data. S.G., W.Y., J.T. and H.P. wrote the manuscript. Y.W., J.T. and H.P. supervised the work. All authors have approved the final version of the manuscript.

## 6 Funding

This work was supported by the National Natural Science Foundation of China (Grant Nos. 51933002 and 51873041) and the National Key R&D Program of China (Grant No. 2016YFC1100300).

## 7 Acknowledgments

We thank Yongbin Cao, Ruihong Xie, Xuechun Zhang and Jingbo Lin at Fudan for helpful discussions.

## 8 Conflict of Interest Statement

The authors declare that the research was conducted in the absence of any commercial or financial relationships that could be construed as a potential conflict of interest.

