## Supplementary material for "Dihydroartemisinin-loaded Magnetic Nanoparticles for Enhanced Chemodynamic Therapy": jing tang

**Supplementary Table S1.** The DLS data of MNP and drug-loaded MNP.

| **Samples** | **Size (nm)** | **PDI** |
|---|---|---|
| **MNP** | 200 | 0.013 |
| **MNP-ART** | 212 | 0.065 |
| **MNP-DHA** | 204 | 0.026 |
| **MNP-AS** | 204 | 0.092 |

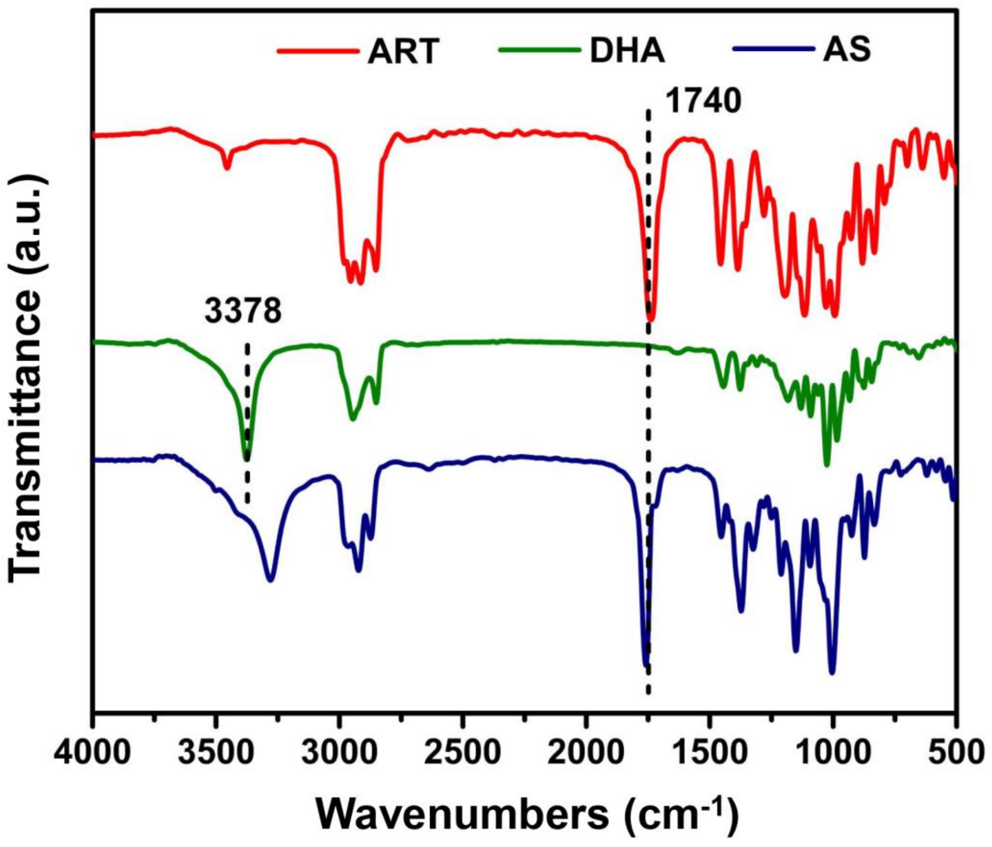


**Supplementary Figure S1.** FT-IR spectra of ART, DHA, and AS.

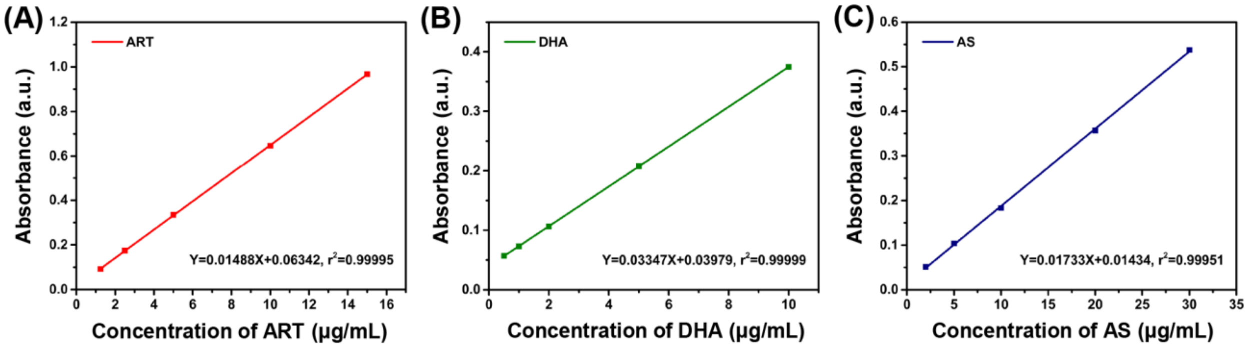


**Supplementary Figure S2.** Standard curves of **(A)** ART, **(B)** DHA, and **(C)** AS concentration.

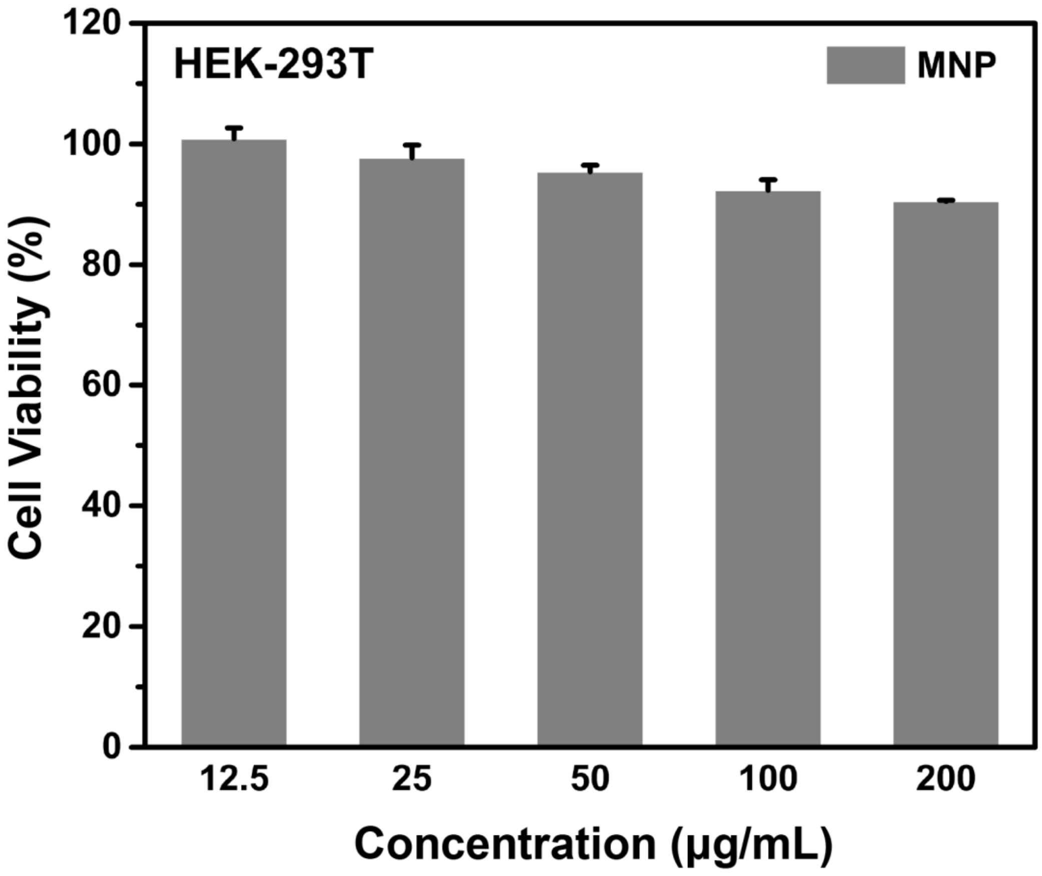


**Supplementary Figure S3.** Cell viability of HEK-293T cells after 24 h incubation with MNP suspensions at different concentrations.
